## Supplemental Information for "Octadecanoids as emerging lipid mediators in cnidarian-dinoflagellate symbiosis"

### Table of contents

**Figure S1.** Cell density curves of *Breviolum minutum* and *Durusdinium trenchii* monocultures….3

**Figure S2:** Sequence alignment of new Symbiodiniaceae LOXes with other 13*S* (At LOX3) and two 9*S* (At LOX1 and At LOX5) LOXes from *Arabidopsis thaliana*…………………………......12

**Table S1:** Quantification of octadecanoids presented as ng/g dry biomass………………………13

**Table S2:** Relative amounts of *R* and *S* enantiomers and calculated **ee** values for octadecanoids in average sample groups…………………………………………………………………………...13

**Table S3**: One-way ANOVA with pairwise Tukey *post hoc* test results…………………………13

**Table S4**: Octadecanoids that differed in the host with a fold-change >2 and $p < 0.05$…………...13

**Table S5**: Octadecanoids that differed in the symbiont with a fold-change >2 and $p < 0.05$……...13

**Table S6:** Species confirmation of symbionts from cultures and symbiotic anemones…………..13

**Table S7:** Description and analytical information about internal standards and new octadecanoids that were added to the original platform…………………………………………………………14

**Table S8:** Genomic stramenopile and alveolate protein references used for Blastp ORF homology……………………………………………………………………………………...15

**Table S9:** GenBank access numbers for phylogeny tree and protein sequency homology………16

**NOTE: The data for Tables S1-S6 and S9 is provided in a separate Excel file**

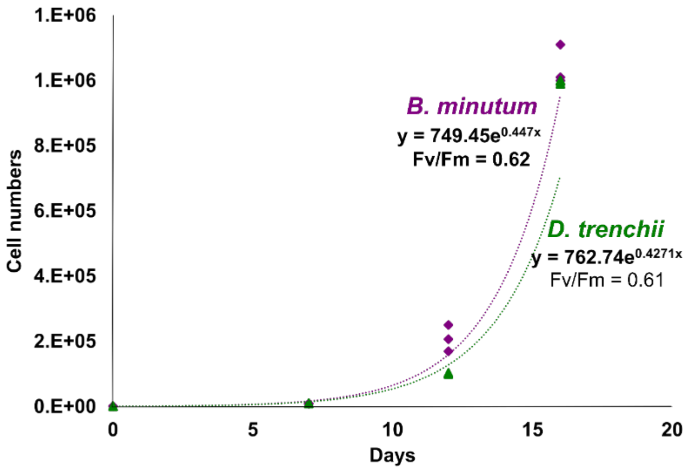


**Figure S1: Cell density curves of *Breviolum minutum* and *Durusdinium trenchii* monocultures (N=3).** Biomass was sampled after 16 days when both species were in exponential growth phase; photosynthetic efficiency ($F_v/F_m$) was also measured at this point. Replicate values were fitted to an exponential curve for the estimation of growth rates.

#### (a) Alignment of new LOX sequences for Symbiodiniaceae with *A. thaliana* LOXes

```
AtLOX3             MALAKELMGYPLITERSSLVSSASHFKKRTQSTQFSINPFDRRPRKTKSGVVAAISEDLV   60
AtLOX1             ------------------------------------------------------MF----   2
AtLOX5             -----------------------------------------------------MIHTDIA   7
Dt_CCMP2556_1      ------------------------------------------------------------   0
Dt_CCMP2556_2      ------------------------------------------------------------   0
Bm_Mf_1.05b.01     ------------------------------------------------------------   0

AtLOX3             KTLRFSTTTGDR--KSEEEEKAAVKFKVRAVVTVRNKNKEDLKETLVKHLDAFADKIGRN   118
AtLOX1             ---------GELRDLLTGGGNETTTKKVKGTVVLMKKNVLDFNDFNASFLDRLHEFLGNK   53
AtLOX5             EILCVKPKTTKKTKTMEEDVKKTTTMKIEGEVVVMKKNLLDFKDVMASLLDRVNELLGRR   67
Dt_CCMP2556_1      ------------------------------------------------------------   0
Dt_CCMP2556_2      ------------------------------------------------------------   0
Bm_Mf_1.05b.01     ------------------------------------------------------------   0

AtLOX3             IVLELISTQLDPKT--KLPKKSNAAVLKDWSKKSKT--KAERVHYTAEFTVDAAFGSPGA   174
AtLOX1             ITLRLVSSDVTDSENGSKGKLGKAAHLEDWITTIT-SLTAGESAFKVTFDYETDFGYPGA   112
AtLOX5             VSLHLISSHQPDPANEKRGRLGKAAHLEKWVTKIKTSVTAEETAFGVTFDWDESMGPPAA   127
Dt_CCMP2556_1      ------------------------------------------------------------   0
Dt_CCMP2556_2      ------------------------------------------------------------   0
Bm_Mf_1.05b.01     ------------------------------------------------------------   0

AtLOX3             ITVMNKHQKEFFLESITIEGFA-----LGPVHFPCNSWVQSQKDHPDKRIFFTNQPYLPN   229
AtLOX1             FLIRNSHFSEFLLKSLTLEDVP----GHGRVHYICNSWIYPAKHYTTDRVFFSNKTYLPH   168
AtLOX5             FVIKNHHHSQFYLKSLTLRGFPDGEGGATAIHFICNSWIYPNHRYRSDRVFFSNKAYLPS   187
Dt_CCMP2556_1      ------------------------------------------------------------   0
Dt_CCMP2556_2      ------------------------------------------------------------   0
Bm_Mf_1.05b.01     ------------------------------------------------------------   0

AtLOX3             ETPSGLRVLREKELKNLRGDG-SGVRKLSDRIYDFDVYNDLGNPDKSSELSRPKLGGK-E   287
AtLOX1             ETPATLLKYREEELVSLRGTG-EGELKEWDRVYDYAYYNDLGVPPKNP---RPVLGGTQE   224
AtLOX5             ETPELIKELREEELKNLRGNEKGGEFKEWDRVYDYAYYNDLGAPDKGPDSVRPVLGGSPE   247
Dt_CCMP2556_1      ------------------------------------------------------------   0
Dt_CCMP2556_2      ------------------------------------------------------------   0
Bm_Mf_1.05b.01     ------------------------------------------------------------   0
```

```
AtLOX3          VPYPRRCRTGRQSTVSDKDAESRVE--KPLPMYVPRDEQFEESKQDTFAAGRLKAVLHHL  345
AtLOX1          YPYPRRGRTGRKPTKEDPQTESRLPITSSLDIYVPRDERFGHLKMSDFLAYALKAIAQFI  284
AtLOX5          LPYPRRGKTGRKSTKSDPKSESRLALL-NLNIYVPRDERFSHVKFSDFLAYALKSVTQVL  306
Dt_CCMP2556_1   ------------------------------------------------------------  0
Dt_CCMP2556_2   ------------------------------------------------------------  0
Bm_Mf_1.05b.01  ------------------------------------------------------------  0

AtLOX3          IPSLKASI--VAEDFADFGEIDRLYKEGLLLKLGFQDDIFKKFPLPKVVVDTLQESTKGL  403
AtLOX1          QPALEAVFDDTPKEFDSFEDVLKIYEEGIDLPNQALIDSIVKNIPLEMLKEIFRTDGQKF  344
AtLOX5          VPEIASVCDKTINEFDSFEDVFHLYDGSIKLANGHTISKLRDVIPWEMFRELVRNDGERF  366
Dt_CCMP2556_1   ------------------------------------------------------------  0
Dt_CCMP2556_2   ------------------------------------------------------------  0
Bm_Mf_1.05b.01  ------------------------------------------------------------  0

AtLOX3          LKYDTPKILSKDKNAWLRDDEFARQAIAGINPVNIERVKTFPPVSNLDPKIYGPQHSALT  463
AtLOX1          LKFPVPQVIKEDKTAWRTDEEFAREMLAGLNPVVIQLLKEFPPKSKLDSESYGNQNSTIT  404
AtLOX5          LKYPLPDILKESRSAWRTDEEFAREMLAGLNPVVISRLQEFPPKSCLDSAKYGNQHSSIR  426
Dt_CCMP2556_1   ------------------------------------------------------------  0
Dt_CCMP2556_2   ------------------------------------------------------------  0
Bm_Mf_1.05b.01  ------------------------------------------------------------  0

AtLOX3          DDHIIGHLDGFSVQQALEENRLYMLDYHDIFLPFLDRINALDGRKAYATRTIFFLTRLGT  523
AtLOX1          KSHIEHNLDGLTVEEALEKERLFILDHHDTLMPYLGRVNTTT-TKTYASRTLLFLKDDGT  463
AtLOX5          TEHIESNMNGLNVQEALEQNKLYILDHHDALMPYLTRINSTN-TKTYATRTLLLLQADGT  485
Dt_CCMP2556_1   ----------------------------------MPRSPISQGNDVYFDKDFR------V  20
Dt_CCMP2556_2   ----------------------------------MPRSPISQGNDAYFDKDFR------V  20
Bm_Mf_1.05b.01  ------------------------------------------------------------  0

AtLOX3          LKPVAIELSLPPHGPKH--R---SKRVLTPPVDATSNWMWQLAKAHVSSNDAGVHQLVNH  578
AtLOX1          LKPLVIELSLPHPNGDKFGA---VSEVYTPGE-GVYDSLWQLAKAFVGVNDSGNHQLISH  519
AtLOX5          LKPLAIELSLPHAQGESYGS---VSKVFTPAEKGVEGSVWQLAKAYAAVNDSGYHQLISH  542
Dt_CCMP2556_1   VK------IVRFENGEVWEDQK-ITTFLPDG-----SRDWEYAKFCFRCSLFTLVTLVDH  68
Dt_CCMP2556_2   VK------IVRFENGEVWEDQK-ITTFLPDG-----SRDWEYAKFCFRCSLFTLVTLVDH  68
Bm_Mf_1.05b.01  -M------IIDQGRGPLVEDFEQKSLVVKPQ-----DDDWERSKFRFRSSLFSLVTLVDH  48
```

```
AtLOX3            WLRTHACLEPF-ILAAHRQLSAMHPIFKLLDPHMRYTLEINALARQSLISADGVIEGGFT   637
AtLOX1            WMQTHASIEPF-VIATNRQLSVLHPVFKLLEPHFRDTMNINALARQILINGGGIFEITVF   578
AtLOX5            WLQTHAVIEPF-IIASNRQLSVVHPIHKLLHPHFRDTMNINALARHVLINSDGVLERTVF   601
Dt_CCMP2556_1     LYGTHLQLANVGVQAMREQLSVDHPVRRFLVPFSYGSININDLARTTLVTRDSWLPRAVA   128
Dt_CCMP2556_2     LYGTHLQLANVGVQAMREQLSVDHPVRRFLVPFSYGSININDLARTTLVTRDSWLPRAVA   128
Bm_Mf_1.05b.01    LYFLHLQLANFFVTSLREQMAEYHPIRRFLTPFTYYTISVNDNAAKNLVAQRSMGPRCFA   108

AtLOX3             AGAYGMEMSAAAYKSSWRFDMEGLPADLI------------RRGMAIPDATQPHGLKLLI   685
AtLOX1             PSKYAMEMSSFIYKNHWTFPDQALPAELK------------KRGMAVEDPEAPHGLRLRI   626
AtLOX5             PSRYAMEMSSSIYK-NWVFTEQALPKDLL------------KRGVAVEDPNSDNGVKLLI   648
Dt_CCMP2556_1      LDDEGLQL-------AWASAFQILPPDYI--TNESDPIKMLESLFDREAQIEKKRSEGL-   178
Dt_CCMP2556_2      LDDEGLQL-------AWASAFQILPPDYI--TNESDPIKMLESLFDREAQIEKKRSEGL-   178
Bm_Mf_1.05b.01     LTDKGFNL-------AFAAAPHLQVWGYEVPAEEGGPFLNLKNYFA------WKRAKG--   153

AtLOX3             EDYPYANDGLLLWSAIQTWVRTYVERYYPNPNL----IKTDSELQSWYSESIN------V   735
AtLOX1             KDYPYAVDGLEVWYAIESWVRDYIFLFYKIEED----IQTDTELQAWWKEVRE------E   676
AtLOX5             EDYPFAVDGLEIWSAIKTWVTEYCTFYYNNDKT----VQTDTEIQSWWTELRT------K   698
Dt_CCMP2556_1      -FTAYYKQALQYWKILHGFVSSYLDHYYGTGAKGDLAMAADQELKLFILQAINLVQTLAS   237
Dt_CCMP2556_2      -FTAYYKQALQYWKILHGFVSSYLDHYYGTGAKGDLAMAADQELKLFILQAINLVQTLAS   237
Bm_Mf_1.05b.01     --------VACFWLI---------------------------------------------   160

AtLOX3             GHADLRDADWWPELSTVDDLVSIL---TTLIWLASAQHAALNFGQYPYGGYVPNRPPLMR   792
AtLOX1             GHGDKKSEPWWPKMQTREELVESC---TIIIWVASALHAAVNFGQYPVAGYLPNRPTISR   733
AtLOX5             GHGDKRHESWWPSMQTRDDLIETC---TIIIWIASALHAAVNFGQYPYAGFLPNRPTVSR   755
Dt_CCMP2556_1      PLTGHRVNELWPHLSDARKRRIMTNFITRFCELSTAGHEQVGDVQ----AY---------   284
Dt_CCMP2556_2      PLTGHRVNELWPHLSDARKRRIMTNFITRFCELSTAGHEQVGDVQ----AY---------   284
Bm_Mf_1.05b.01     ------------------------------------------------------------   160

AtLOX3             RLIPDESDPEYASFISHPEKYYFSSMPSLAQTSKFMAVVDTLSTHSPDEEYIGERQQPSI   852
AtLOX1             QYMPKENTPEFEELEKNPDKVFLKTITAQLQTLLGISLIEILSTHSSDEVYLGQRDSK-E   792
AtLOX5             RFMPEPGTDEYAELEEDADVAFLKTITPQLQTLLGISIIEILSMHSTDEIYLGQRDSP-N   814
Dt_CCMP2556_1      -----AQDPSFCSFSWPKSLRREGALVAPKEVGLGVALIMALT--STPMPRLLVREPTDD   337
Dt_CCMP2556_2      -----AQDPSFCSFSWPKSLRREGALVAPKEVGLGVALIMALT--STPMPRLLVREPTDD   337
Bm_Mf_1.05b.01     ------------------------------------------------------------   160
```

```
AtLOX3              WTG---------DAEIVEAFYGFAAEIGRIEKEIEKRNADPDRRNRCGAGVLPYELLVPS   903
AtLOX1              WAA---------EKEALEAFEKFGEKVKEIEKNIDERNDDETLKNRTGLVKMPYTLLFPS   843
AtLOX5              WTA---------DDEPLEAFKRFGKELELIENNIIRRNNDKRFKNRTGPVNIPYTLLYPN   865
Dt_CCMP2556_1       WSHLFPGASEADRSKLNSIFERFQSELQTFSKEC--------------------------   371
Dt_CCMP2556_2       WSHLFPGASEADRSKLNSIFERFQSELQTFSKECD-------------------------   372
Bm_Mf_1.05b.01      ------------------------------------------------------------   160

AtLOX3              -----SEPGVTCRGVPNSVSI    919
AtLOX1              -----SEGGVTGRGIPNSVSI    859
AtLOX5              TTDYTREGGITGKGIPNSVSI    886
Dt_CCMP2556_1       ---------------------    371
Dt_CCMP2556_2       ---------------------    372
Bm_Mf_1.05b.01      ---------------------    160
```

#### (b) Alignment of GenBank complete sequences with *A. thaliana* LOX sequences

```
Dt_CCMP2556_1      ------------------------------------------------------------   0
Dt_CCMP2556_2      ------------------------------------------------------------   0
CAK8999115.1       ------------------------------------------------------------   0
CAK9070246.1       ------------------------------------------------------------   0
AtLOX3             MALAKELMGYPLITERSSLVSSASHFKKRTQSTQFSINPFDRRPRKTKSGVVAAISEDLV   60
AtLOX1             ------------------------------------------------------MF----   2
AtLOX5             -----------------------------------------------------MIHTDIA   7

Dt_CCMP2556_1      ------------------------------------------------------------   0
Dt_CCMP2556_2      ------------------------------------------------------------   0
CAK8999115.1       ------------------------------------------------------------   0
CAK9070246.1       ------------------------------------------------------------   0
AtLOX3             KTLRFSTTTGDR--KSEEEEKAAVKFKVRAVVTVRNKNKEDLKETLVKHLDAFADKIGRN   118
AtLOX1             ---------GELRDLLTGGGNETTTKKVKGTVVLMKKNVLDFNDFNASFLDRLHEFLGNK   53
AtLOX5             EILCVKPKTTKKTKTMEEDVKKTTTMKIEGEVVVMKKNLLDFKDVMASLLDRVNELLGRR   67

Dt_CCMP2556_1      ------------------------------------------------------------   0
Dt_CCMP2556_2      ------------------------------------------------------------   0
CAK8999115.1       ------------------------------------------------------------   0
CAK9070246.1       ------------------------------------------------------------   0
AtLOX3             IVLELISTQLDPKT--KLPKKSNAAVLKDWSKKSKT--KAERVHYTAEFTVDAAFGSPGA   174
AtLOX1             ITLRLVSSDVTDSENGSKGKLGKAAHLEDWITTIT-SLTAGESAFKVTFDYETDFGYPGA   112
AtLOX5             VSLHLISSHQPDPANEKRGRLGKAAHLEKWVTKIKTSVTAEETAFGVTFDWDESMGPPAA   127

Dt_CCMP2556_1      ------------------------------------------------------------   0
Dt_CCMP2556_2      ------------------------------------------------------------   0
CAK8999115.1       ------------------------------------------------------------   0
CAK9070246.1       ------------------------------------------------------------   0
AtLOX3             ITVMNKHQKEFFLESITIEGFA-----LGPVHFPCNSWVQSQKDHPDKRIFFTNQPYLPN   229
AtLOX1             FLIRNSHFSEFLLKSLTLEDVP----GHGRVHYICNSWIYPAKHYTTDRVFFSNKTYLPH   168
AtLOX5             FVIKNHHHSQFYLKSLTLRGFPDGEGGATAIHFICNSWIYPNHRYRSDRVFFSNKAYLPS   187

Dt_CCMP2556_1      ------------------------------------------------------------   0
Dt_CCMP2556_2      ------------------------------------------------------------   0
```

```
CAK8999115.1       ------------------------------------------------MEQSPLIGESLY   12
CAK9070246.1       ------------------------------------------------MEQSPLIGESLY   12
AtLOX3             ETPSGLRVLREKELKNLRGDG-SGVRKLSDRIYDFDVYNDLGNPDKSSELSRPKLGGK-E   287
AtLOX1             ETPATLLKYREEELVSLRGTG-EGELKEWDRVYDYAYYNDLGVPPKNP---RPVLGGTQE   224
AtLOX5             ETPELIKELREEELKNLRGNEKGGEFKEWDRVYDYAYYNDLGAPDKGPDSVRPVLGGSPE   247

Dt_CCMP2556_1      ------------------------------------------------------------   0
Dt_CCMP2556_2      ------------------------------------------------------------   0
CAK8999115.1       A--------------------------PSWPVVQPPEPSFYSLGCCQTCLRIPAAFGCGG   46
CAK9070246.1       A--------------------------PSWPVVQPPEPSFYSLGCCQTCLRIPAAFGCGG   46
AtLOX3             VPYPRRCRTGRQSTVSDKDAESRVE--KPLPMYVPRDEQFEESKQD--------------   331
AtLOX1             YPYPRRGRTGRKPTKEDPQTESRLPITSSLDIYVPRDERFGHLKMS--------------   270
AtLOX5             LPYPRRGKTGRKSTKSDPKSESRLALL-NLNIYVPRDERFSHVKFS--------------   292

Dt_CCMP2556_1      ------------------------------------------------------------   0
Dt_CCMP2556_2      ------------------------------------------------------------   0
CAK8999115.1       CRICCYSILAVL-------------------LFALLLGFYPIVVALVLVLLLLLGLPISY   87
CAK9070246.1       CRICCYSILAVL-------------------LFALLLGFYPIVVALVLVLLLLLGLPISY   87
AtLOX3             -TFAAGRLKAVLHHLIPSLKASI--VAEDFADFGEIDRLYKEG------LLLKLGFQD--   380
AtLOX1             -DFLAYALKAIAQFIQPALEAVFDDTPKEFDSFEDVLKIYEEG------IDLPNQALI--   321
AtLOX5             -DFLAYALKSVTQVLVPEIASVCDKTINEFDSFEDVFHLYDGS------IKLANGHTI--   343

Dt_CCMP2556_1      ------------------------------------------------------------   0
Dt_CCMP2556_2      ------------------------------------------------------------   0
CAK8999115.1       AMIYLLFLNPPPELAWKIAKLTQLLRLAFYQFPKPA-------EWRVDQLAPKGPLETGL   140
CAK9070246.1       AMIYLLFLNPPPELAWKIAKLTQLLRLAFYQFPKPA-------EWRVDQLAPKGPLETGL   140
AtLOX3             ---DIFKKFPLPKVVVDT---LQESTKGLLKYDTPKILSKDKNAWLRDDEFARQAI-AGI   433
AtLOX1             ---DSIVKNIPLEMLKEI---FRTDGQKFLKFPVPQVIKEDKTAWRTDEEFAREML-AGL   374
AtLOX5             ---SKLRDVIPWEMFREL---VRNDGERFLKYPLPDILKESRSAWRTDEEFAREML-AGL   396

Dt_CCMP2556_1      ------------------------------------------------------------   0
Dt_CCMP2556_2      ------------------------------------------------------------   0
CAK8999115.1       LQYAIAIRVPFLQQDELYAGGLFERFLRSSPSTEFVTSSVFST----LP----QMELKDM   192
CAK9070246.1       SQYAIAIRVPFLQQDELYAGGLFERFLRSSPSTEFVTSSVFST----LP----QMELKDM   192
AtLOX3             NPVNIERV-------------------KTFPPVSNLDPKIYGPQHSALTDDHIIGHLDGF   474
```

```
AtLOX1             NPVVIQLL-------------------KEFPPKSKLDSESYGNQNSTITKSHIEHNLDGL    415
AtLOX5             NPVVISRL-------------------QEFPPKSCLDSAKYGNQHSSIRTEHIESNMNGL    437

Dt_CCMP2556_1      ------------------------------------------------------------    0
Dt_CCMP2556_2      ------------------------------------------------------------    0
CAK8999115.1       SLFKEGENPVEYVMGVVQDIYPRINQEWTDKTSDRALTHLCLHGLGAHRLERADSTHPGC    252
CAK9070246.1       SLFKEGENPVEYVMGVVQDIYPRINQEWTDKTSDRALTHLCLHGLGAHRLERADSTHPGC    252
AtLOX3             SVQQALEENRLYMLD--------------------------YHDIFLPFLDRINALDGRK    508
AtLOX1             TVEEALEKERLFILD--------------------------HHDTLMPYLGRVNTTT-TK    448
AtLOX5             NVQEALEQNKLYILD--------------------------HHDALMPYLTRINSTN-TK    470

Dt_CCMP2556_1      -------------MPRSPISQGNDVYFDKDFRVVKIVR------FENGEVWED-QKITTF    40
Dt_CCMP2556_2      -------------MPRSPISQGNDAYFDKDFRVVKIVR------FENGEVWED-QKITTF    40
CAK8999115.1       SYVVRTNQLSTLPVREGYETYGNDVYFDKDFRVVKIVR------FENGEVWED-QKITTF    305
CAK9070246.1       SYVVRTNQLSTLPVREGYETYGNDAYFDKDFRVVKIVR------FENGEVWED-QKITTF    305
AtLOX3             AYATRTI------------------FFLTRLGTLKPVAIELSLPPHGPKH--RSKRVLTP    548
AtLOX1             TYASRTL------------------LFLKDDGTLKPLVIELSLPHPNGDKFGAVSEVYTP    490
AtLOX5             TYATRTL------------------LLLQADGTLKPLAIELSLPHAQGESYGSVSKVFTP    512

Dt_CCMP2556_1      LPDGS--RDWEYAKFCFRCSLFTLVTLVDHLYGTHLQLANVGVQAMREQLSVDHPVRRFL    98
Dt_CCMP2556_2      LPDGS--RDWEYAKFCFRCSLFTLVTLVDHLYGTHLQLANVGVQAMREQLSVDHPVRRFL    98
CAK8999115.1       LPDGS--RDWEYAKFCFRCSLFTLVTLVDHLYGTHLQLANVGVQAMREQLSVDHPVRRFL    363
CAK9070246.1       LPDGS--RDWEYAKFCFRCSLFTLVTLVDHLYGTHLQLANVGVQAMREQLSVDHPVRRFL    363
AtLOX3             PVDATSNWMWQLAKAHVSSNDAGVHQLVNHWLRTHACLEPF-ILAAHRQLSAMHPIFKLL    607
AtLOX1             GE-GVYDSLWQLAKAFVGVNDSGNHQLISHWMQTHASIEPF-VIATNRQLSVLHPVFKLL    548
AtLOX5             AEKGVEGSVWQLAKAYAAVNDSGYHQLISHWLQTHAVIEPF-IIASNRQLSVVHPIHKLL    571

Dt_CCMP2556_1      VPFSYGSININDLARTTLVTRDSWLPRAVALDDEGLQ-------LAWASAFQILPPDYIT    151
Dt_CCMP2556_2      VPFSYGSININDLARTTLVTRDSWLPRAVALDDEGLQ-------LAWASAFQILPPDYIT    151
CAK8999115.1       VPFSYGSININDLARTTLVTRDSWLPRAVALDDEGLQ-------LAWASAFQILPPDYIT    416
CAK9070246.1       VPFSYGSININDLARTTLVTRDSWLPRAVALDDEGLQ-------LAWASAFQILPPDYIT    416
AtLOX3             DPHMRYTLEINALARQSLISADGVIEGGFTAGAYGMEMSAAAYKSSWRFDMEGLPADLIR    667
AtLOX1             EPHFRDTMNINALARQILINGGGIFEITVFPSKYAMEMSSFIYKNHWTFPDQALPAELKK    608
AtLOX5             HPHFRDTMNINALARHVLINSDGVLERTVFPSRYAMEMSSSIYK-NWVFTEQALPKDLLK    630
```

```
Dt_CCMP2556_1     NESDPIKMLESLFDREAQIEKKR-SEGL-----FTAYYKQALQYWKILHGFVSSYLDHYY   205
Dt_CCMP2556_2     NESDPIKMLESLFDREAQIEKKR-SEGL-----FTAYYKQALQYWKILHGFVSSYLDHYY   205
CAK8999115.1      NESDPIKMLESLFDREAQIEKKR-SEGL-----FTAYYKQALQYWKILHGFVSSYLDHYY   470
CAK9070246.1      NESDPIKMLESLFDREAQIEKKR-SEGL-----FTAYYKQALQYWKILHGFVSSYLDHYY   470
AtLOX3            RGM--------------AIPDATQPHGLKLLIEDYPYANDGLLLWSAIQTWVRTYVERYY   713
AtLOX1            RGM--------------AVEDPEAPHGLRLRIKDYPYAVDGLEVWYAIESWVRDYIFLFY   654
AtLOX5            RGV--------------AVEDPNSDNGVKLLIEDYPFAVDGLEIWSAIKTWVTEYCTFYY   676

Dt_CCMP2556_1     GTGAKGDLAMAADQELKLFILQAINLVQTLASPLTGHRVNELWPHLSDARKRRIMTNFIT   265
Dt_CCMP2556_2     GTGAKGDLAMAADQELKLFILQAINLVQTLASPLTGHRVNELWPHLSDARKRRIMTNFIT   265
CAK8999115.1      GTGAKGDLAMAADQELKLFILQAINLVQTLASPLTGHRVNELWPHLSDARKRRIMTNFIT   530
CAK9070246.1      GTGAKGDLAMAADQELKLFILQAINLVQTLASPLTGHRVNELWPHLSDARKRRIMTNFIT   530
AtLOX3            PNPNL----IKTDSELQSWYSESIN------VGHADLRDADWWPELSTVDDLVSIL---T   760
AtLOX1            KIEED----IQTDTELQAWWKEVRE------EGHGDKKSEPWWPKMQTREELVESC---T   701
AtLOX5            NNDKT----VQTDTEIQSWWTELRT------KGHGDKRHESWWPSMQTRDDLIETC---T   723

Dt_CCMP2556_1     RFCELSTAGHEQVGDVQ----AY--------------AQDPSFCSFSWPKSLRREGALVA   307
Dt_CCMP2556_2     RFCELSTAGHEQVGDVQ----AY--------------AQDPSFCSFSWPKSLRREGALVA   307
CAK8999115.1      RFCELSTAGHEQVGDVQ----AY--------------AQDPSFCSFSWPKSLRREGALVA   572
CAK9070246.1      RFCELSTAGHEQVGDVQ----AY--------------AQDPSFCSFSWPKSLRREGALVA   572
AtLOX3            TLIWLASAQHAALNFGQYPYGGYVPNRPPLMRRLIPDESDPEYASFISHPEKYYFSSMPS   820
AtLOX1            IIIWVASALHAAVNFGQYPVAGYLPNRPTISRQYMPKENTPEFEELEKNPDKVFLKTITA   761
AtLOX5            IIIWIASALHAAVNFGQYPYAGFLPNRPTVSRRFMPEPGTDEYAELEEDADVAFLKTITP   783

Dt_CCMP2556_1     PKEVGLGVALIMALT--STPMPRLLVREPTDDWSHLFPGASEADRSKLNSIFERFQSELQ   365
Dt_CCMP2556_2     PKEVGLGVALIMALT--STPMPRLLVREPTDDWSHLFPGASEADRSKLNSIFERFQSELQ   365
CAK8999115.1      PKEVGLGVALIMALT--STPMPRLLVREPTDDWSHLFPGASEADRSKLNSIFERFQSELQ   630
CAK9070246.1      PKEVGLGVALIMALT--STPMPRLLVREPTDDWSHLFPGASEADRSKLNSIFERFQSELQ   630
AtLOX3            LAQTSKFMAVVDTLSTHSPDEEYIGERQQPSIWTG---------DAEIVEAFYGFAAEIG   871
AtLOX1            QLQTLLGISLIEILSTHSSDEVYLGQRDSK-EWAA---------EKEALEAFEKFGEKVK   811
AtLOX5            QLQTLLGISIIEILSMHSTDEIYLGQRDSP-NWTA---------DDEPLEAFKRFGKELE   833

Dt_CCMP2556_1     TFSKEC------------------------------------------------------   371
Dt_CCMP2556_2     TFSKECD-----------------------------------------------------   372
CAK8999115.1      TFSKECDDYNAEAKNRAFPNDFGLWVFN-----PK-------------YLETSVSI      668
```

```
CAK9070246.1        TFSKECDDYNAEAKNRAFPNDFGLWVFN-----PK-------------YLETSVSI        668

AtLOX3              RIEKEIEKRNADPDR---RNRCGAGVLPYELLVPS-----SEPGVTCRGVPNSVSI        919

AtLOX1              EIEKNIDERNDDETL---KNRTGLVKMPYTLLFPS-----SEGGVTGRGIPNSVSI        859

AtLOX5              LIENNIIRRNNDKRF---KNRTGPVNIPYTLLYPNTTDYTREGGITGKGIPNSVSI        886
```

**Figure S2: Sequence alignment of new Symbiodiniaceae LOXes with other 13*S* (At LOX3) and two 9*S* (At LOX1 and At LOX5) LOXes from *Arabidopsis thaliana*.** (**a**) According to the alignment, the *Durisdinium trenchii* (Dt) sequences are most likely 9*S*-LOXes and the *Breviolum minutum* (Bm) sequence is a 13*S*-LOX, defined by the Hornung (yellow/green) and Coffa (blue) sites (Wasternack & Feussner, 2018). The sequences are partial, and the one obtained for *B. minutum* was not found in any publicly available dataset. (**b**) The complete sequences for *D. trenchii* were found in GenBank (CAK8999115.1 and CAK9070246.1) annotated as "unnamed proteins" and again aligned with the same *A. thaliana* LOXes. *See Excel file in electronic version.*

**Table S1:** Quantification of octadecanoids presented as ng/g dry biomass. *See Excel file in electronic version.*

**Table S2:** Relative amounts of *R* and *S* enantiomers and calculated **ee** values for octadecanoids in average sample groups. Values suggesting enzymatic biosynthesis are highlighted. *See Excel file in electronic version.*

**Table S3**: One-way ANOVA with pairwise Tukey *post hoc* test results. Only compounds that were statistically significant ($p < 0.05$) with FDR are listed. *See Excel file in electronic version.*

**Table S4**: Octadecanoids that differed in the host with a fold-change >2 and $p < 0.05$. Pairwise comparisons were between symbiotic hosts, colonized with *Breviolum minutum* (Host_B) and *Durusdinium trenchii* (Host_D), and *versus* the aposymbiotic host (Host_Apo). *See Excel file in electronic version.*

**Table S5**: Octadecanoids that differed in the symbiont with a fold-change >2 and $p < 0.05$. Pairwise comparisons were between the symbionts *Breviolum minutum* (B_symbiotic) and *Durusdinium trenchii* (D_symbiotic) when in symbiosis, and when isolated in culture (B_cultured and D_cultured, respectively). *See Excel file in electronic version.*

**Table S6:** Species confirmation of symbionts from cultures and symbiotic anemones. *See Excel file in electronic version*

**Table S7:** Internal standards and additional octadecanoids that were included in the current work that are not described in the original method of Quaranta *et al*. (2022).

| Internal standard full name | Abbreviation | Parent PUFA | Detected | RT (min) | Source |
|---|---|---|---|---|---|
| *cis*-12,13-epoxy-octadecenoic acid-d4 | *cis*-12,13-EpoME-d4 | LA | N/A | 4.18 | In-house |
| 13-oxo-octadecadienoic acid-d3 | 13-oxo-ODE-d3 | LA | N/A | 4.15 | Cayman |
| 9(*S*)-hydroxy-octadecadienoic acid-d4 | 9(*S*)-HODE-d4 | LA | N/A | 6.93 | Cayman |
| 13(*S*)-hydroxy-octadecadienoic acid-d4 | 13(*S*)-HODE-d4 | LA | N/A | 8.14 | Cayman |
| *threo*-9,10-dihydroxy-octadecenoic acid-d4 | *threo*-9,10-DiHOME-d4 | LA | N/A | 8.48 | Cayman |
| *threo*-12,13-dihydroxy-octadecenoic acid-d4 | threo-12,13-DiHOME-d4 | LA | N/A | 7.42 | Cayman |
| 9(*S*),12(*S*),13(*S*)-13C3-trihydroxy-octadecenoic acid | 9(*S*),12(*S*),13(*S*)-13C3-TriHOME | LA | N/A | 11.14 | In house |
| **New octadecanoid full name** | **Abbreviation** | **Parent PUFA** | **Detected** | **RT (min)** | **Source** |
| 10-oxo-decanoic acid | 10-oxo-ODA | OA | YES | 3.56 | In house |
| 13-oxo-decanoic acid' | 13-oxo-ODA | OA | YES | 3.59 | In house |
| 13(*R*)-9(*Z*),15(*Z*)-hydroxy-octadecadienoic acid | 9(*Z*),15(*Z*)-13(*R*)-HODE | ALA | YES | 4.67 | In house |
| 13(S)-9(Z),15(Z)-hydroxy-octadecadienoic acid | 9(Z),15(Z)-13(*S*)-HODE | ALA | YES | 4.94 | In house |
| 10(R)-8(E),12(Z)-hydroxy-octadecadienoic acid | 8(E),12(Z)-10(*R*)-HODE | ALA | YES | 4.34 | In house |
| 10(S)-8(E),12(Z)-hydroxy-octadecadienoic acid | 8(E),12(Z)-10(*S*)-HODE | ALA | YES | 4.75 | In house |
| 12(R)-9(Z),13(E)-hydroxy-octadecadienoic acid | 9(Z),13(E)-12(*R*)-HODE | ALA | YES | 4.65 | In house |
| 12(S)-9(Z),13(E)-hydroxy-octadecadienoic acid | 9(Z),13(E)-12(*S*)-HODE | ALA | YES | 4.83 | In house |
| 9(R)-6(Z),10(E),12(Z)-hydroxy-octadecatrienoic acid | 9(*R*)-HOTrE-ɣ | GLA | YES | 4.07 | In house |
| 9(S)-6(Z),10(E),12(Z)-hydroxy-octadecatrienoic acid | 9(*S*)-HOTrE-ɣ | GLA | YES | 4.54 | In house |
| 10(R)-6(Z),8(E),12(Z)-hydroxy-octadecatrienoic acid | 10(*R*)-HOTrE-ɣ | GLA | YES | 4.07 | In house |
| 10(S)-6(Z),8(E),12(Z)-hydroxy-octadecatrienoic acid | 10(*S*)-HOTrE-ɣ | GLA | YES | 4.40 | In house |
| 12(R,S)-6(Z),9(Z),13(E)-hydroxy-octadecatrienoic acid | 12(*R,S*)-HOTrE-ɣ | GLA | YES | 4.07 | In house |
| 13(R)-6(Z),9(Z),11(E)-hydroxy-octadecatrienoic acid | 13(*R*)-HOTrE-ɣ | GLA | YES | 4.20 | In house |
| 13(S)-6(Z),9(Z),11(E)-hydroxy-octadecatrienoic acid | 13(*S*)-HOTrE-ɣ | GLA | YES | 6.21 | In house |

| | | | | | |
|---|---|---|---|---|---|
| 9(*R*)-hydroxy-octadecatetraenoic acid | 9(*R*)-HOTE | SDA | YES | 4.56 | In house |
| 9(*S*)-hydroxy-octadecatetraenoic acid | 9(*S*)-HOTE | SDA | YES | 5.17 | In house |
| 10(*R*)-hydroxy-octadecatetraenoic acid | 10(*R*)-HOTE | SDA | YES | 4.23 | In house |
| 10(*S*)-hydroxy-octadecatetraenoic acid | 10(*S*)-HOTE | SDA | YES | 4.58 | In house |
| 12(*R*)-hydroxy-octadecatetraenoic acid | 12(*R*)-HOTE | SDA | YES | 4.26 | In house |
| 12(*S*)-hydroxy-octadecatetraenoic acid | 12(*S*)-HOTE | SDA | YES | 4.49 | In house |
| 13(*R*)-hydroxy-octadecatetraenoic acid | 13(*R*)-HOTE | SDA | YES | 4.14 | In house |
| 13(*S*)-hydroxy-octadecatetraenoic acid | 13(*S*)-HOTE | SDA | YES | 4.86 | In house |
| 15(*R,S*)-hydroxy-octadecatetraenoic acid | 15(*R,S*)-HOTE | SDA | YES | 4.38 | In house |
| 16(*R*)-hydroxy-octadecatetraenoic acid | 16(*R*)-HOTE | SDA | YES | 4.25 | In house |
| 16(*S*)-hydroxy-octadecatetraenoic acid | 16(*S*)-HOTE | SDA | YES | 4.67 | In house |
| 18-hydroxy-octadecatetraenoic acid | 18-HOTE | SDA | YES | .64 | In house |

**Table S8:** Genomic stramenopile and alveolate protein references used for Blastp ORF homology filtering of *Breviolum minutum* and *Durusdinium trenchii* transcriptomes. *See electronic version*

| Species | Strain / isolate | WGS project | Assembly Accession | Submission date | Contigs | Encoded proteins |
|---|---|---|---|---|---|---|
| *Amoebophrya* sp. | A120 | CAJINV01 | GCA_905178155.1 | Sep 28, 2021 | 1,147 | 26,350 |
| *Cyanidium caldarium* | DBV 063 E5 | JANCYW01 | GCA_026184775.1 | Nov 15, 2022 | 20 | 4,870 |
| *Ectocarpus* sp. | EcLAC-371 | CAXNAH01 | GCA_964200375.1 | Jul 24, 2024 | 436 | 685 |
| *Leishmania donovani* | - | LDHU3_new | GCA_900635355.2 | Jun 28, 2020 | 36 | 8,463 |
| *Phaeodactylum tricornutum* | CCAP 1055/1 | CCAP1055 | GCA_000150955.2 | Dec 12, 2008 | 179 | 10,392 |
| *Plasmodium falciparum* | 3D7 | PRJNA13173 | GCA_000002765.3 | Apr 7, 2016 | 14 | 5,387 |
| *Thalassiosira pseudonana* | CCMP1335 | AAFD02 | GCA_000149405.2 | Jan 16, 2009 | 115 | 11,672 |

**Table S9:** GenBank access numbers for phylogeny tree and protein sequency homology. *See Excel file in electronic version*
